# Slower-than-exponential viral decay is prevalent and can reshape virus-microbe dynamics

**DOI:** 10.64898/2026.08.27.747580

**Authors:** Akash Arani, Paul Frémont, Emma Wachter, Joshua S. Weitz

**Affiliations:** Department of Biology, University of Maryland, College Park, MD, USA; Department of Physics, University of Maryland, College Park, MD, USA; University of Maryland Institute for Health Computing, North Bethesda, MD, USA

## Abstract

Viral population dynamics are shaped by production and loss. For viruses of microbes, high standing levels of viral abundances are interpreted as evidence of high rates of viral-induced cellular loss and viral production, followed by rapid extracellular viral decay. Here we reassess assumptions of rapid extracellular decay in 17 curated datasets, finding that biphasic decay either fits better or is statistically indistinguishable from exponential decay in approximately half the datasets. In addition to intrinsic heterogeneity in decay rates, biphasic decay at population scales can arise generically through aggregation mechanisms, where single virions decay and viral aggregates are protected. Integrating aggregation-induced biphasic decay into a virus-host model reveals that accounting for aggregation can recapitulate joint observations of high virion abundances and low infection prevalence, without assuming significant levels of uniformly in-efficient infection. Together, our results suggest that durable extracellular virion persistence is environmentally relevant in shaping virus–microbe population dynamics.

## Introduction

Viruses of microbes (VOMs) are ubiquitous across ecosystems, with total densities of virus-like particles often exceeding 10^8^/ml in oceans [1], 10^10^/ml in hypersaline environments [2], 10^8^/g in soil [3], and 10^8^/g in the gut [4]. These high abundances reflect a balance between viral production and loss. Individual virions can lose infectivity through several mechanisms, including damage to the viral genome, aggregation, physical degradation, UV exposure, and thermal destabilization [5, 6, 7, 8, 9]. Conventional virus-host models typically assume that the combined effects of these processes can be represented as rapid extracellular decay with a single characteristic exponential timescale. As a result, high standing VOM abundances are often interpreted as evidence that viruses infect hosts and produce at high rates to balance the rapid exponential loss of free virions (via particle decay and/or loss of infectiousness) [10, 11, 12, 13].

Recent studies however, suggest that VOM decay is not always well described by an exponential curve [14, 15, 16, 17]. Biphasic, multiphasic, and generalized non-exponential decay indicates that a subset of virions may persist substantially longer than expected under a single rapid exponential decay process [9, 17]. Similarly, experimental studies have found that microbial infection levels can remain unexpectedly low even when viruses are highly abundant and outnumber their microbial hosts [18, 19, 20, 21]. Together, these observations challenge the conventional view that high viral abundances necessarily require a balance of rapid extracellular decay, high infection/mortality rates, and high viral production. Instead, they raise the possibility that abundant viral populations may be sustained, at least in part, by durable extracellular persistence rather than continual high turnover. Mechanisms capable of generating persistent virions are already well established in models of viruses of humans (VOHs), where processes such as extracellular aggregation [22, 23], intracellular latent reservoirs [24, 25, 26], and heterogeneity in virion stability [27, 28] have been incorporated into models and shown to substantially alter inferred dynamics [24, 29, 26, 30, 31]. By contrast, analogous mechanisms, while observed in VOMs [9, 14, 32, 33], are rarely incorporated into VOM models, leaving their consequences for interpreting viral abundance and host infection and killing rates largely unresolved.

In this study, we re-examine the conventional assumption of fast, exponential decay of extracellular virions in virus-microbe systems. We first conduct a systematic literature search and identify 17 VOM decay datasets with sufficient temporal resolution to statistically compare the quality of fits of exponential and biphasic decay models, finding that biphasic decay is frequently, though not universally, supported. We then develop a mechanistic model of reversible extracellular virion aggregation, a feature that has been observed repeatedly in VOMs [34, 32, 33], in which virions can enter a temporarily protected aggregated state before returning to the infectious virion pool. This mechanism reproduces biphasic decay dynamics at the population scale, with aggregation-associated parameters that can be partially mapped from subpopulation-based biphasic decay parameters. Finally, we examine how aggregation reshapes virus–host dynamics at the population level. By sequestering virions into a non-infectious aggregate pool, aggregation can facilitate the persistence of high viral abundances with low levels of host infection, thereby decoupling viral standing abundance from realized infection pressure. This framework suggests a continuum between inefficient adsorption across the total viral pool and more efficient adsorption within a smaller infectious subpopulation. Together, our results suggest that accounting for slower-than-exponential extracellular decay may require reassessing conventional estimates of viral-induced mortality rates in ecologically relevant regimes.

## Materials and Methods

### Selecting and extracting data from published studies

To identify studies with viral decay data suitable for our analyses, we conducted a single Web of Science search using the following search query: *“virus decay” OR “viral decay” OR “bacterio-phage decay” OR “phage decay”) AND (“bacteriophage” OR “phage “ OR “cyanophage” OR “marine” OR “algae” OR “plankton” OR “T7” OR “T4” OR “lambda” OR “aquatic”)*. We restricted inclusion to datasets that contained at least five data points, as the biphasic model requires four parameters. We recognize that a higher ratio of data points to parameters would have been preferable, such datasets were uncommon.

Of the 175 studies retrieved through the keyword search, only the 17 papers listed in Table 1 met our stringent criteria. As most of the studies did not provide raw numerical decay data, we used PlotDigitizer to extract data from the published graphs. Extracted data included the individual points and, when available, the associated error bars. When replicate measurements were reported at the same time point, values were averaged prior to analysis.

### Viral Decay Models

We focus on two models of virion decay: monophasic exponential decay and biphasic decay. In the case of monophasic exponential decay, viral load (*V*) decreases over time (*t*) according to:

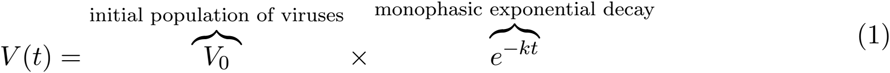

where *k* is the exponential decay constant. In the case of biphasic decay, viral load (*V*) decreases over time (*t*) according to:

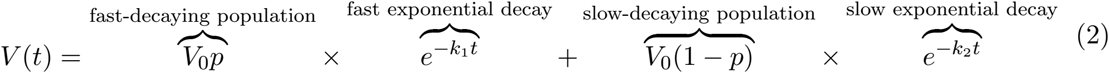

where *p* is the proportion of the population that decays at exponential rate *k*_1_, and 1 *− p* is the proportion of the population that decays at a slower exponential rate *k*_2_. We chose the biphasic model to represent non-exponential dynamics because it has been experimentally observed in both viruses of humans and microbes [9, 23, 31, 24], and because previous studies have shown biphasic decay to be more parsimonious for describing non-exponential decay than triphasic or higher-order multiphasic models [9]. Refer to Table S1 for model parameter and state variable descriptions.

### Virus-Host Models

We use a conventional nonlinear virus-host model [10], which tracks susceptible hosts, infected hosts, and free virions as follows:

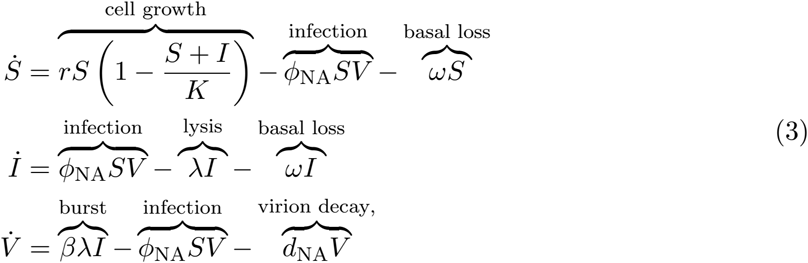

and extended it by incorporating reversible aggregation of free infectious virions into non-infectious aggregates. The resulting aggregation model tracks susceptible hosts, and infected hosts, free virions, and viral aggregates as follows:

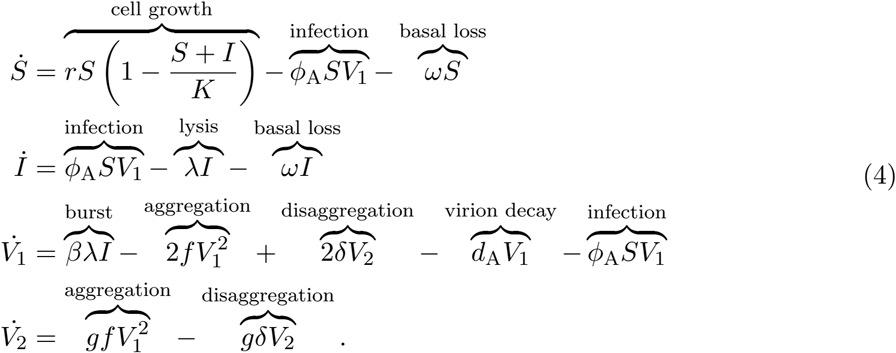

In both models, *S, I*, and *V* denote susceptible cells, infected cells, and viruses, respectively, while the aggregation model distinguishes between single viruses *V*_1_ that decay and are infectious, and viral aggregates *V*_2_ that do not decay and are non-infectious. Here, *f* and *δ* denote the aggregation and disaggregation rates, respectively, and *g* accounts for the measurement method used to quantify aggregates. Typically, we set *g* = 1 as studies of viruses of microbes often use spatially separated measurement methods, including polony, VLP, and PFU assays, which count an aggregate as a single measured viral unit; whereas *g* = 2 is appropriate when genome-based measurements of viral density are used. The extracellular aggregation framework and detailed descriptions of *f*, *δ* and *g* are described in greater detail in *Results: Extracellular virion aggregation as a proposed generic mechanism that can drive non-exponential decay*. The adsorption rate of a virion to a host *ϕ* and the single virion decay rate *d* are different between the models as they represent infection and decay features of the entire virion population in the non-aggregation model, but solely of the infectious single virion (*V*_1_) subpopulation in the aggregation model. Additional parameters that are equivalent in both models include the growth rate of the host cells *r*, cellular carrying capacity *K*, basal cellular loss rate *ω*, lysis rate *λ*, burst size *β*, and the genome measurement parameter *g*.

### Viral Decay Model Fitting and Selection

All models were fit to normalized viral decay data (*V*_0_ = 1) in log_10_ space using nonlinear least-squares optimization implemented with the `lmfit` package [35] in Python 3.0. Model parameters were estimated using the Levenberg–Marquardt algorithm available in the `lmfit` package, a gradient based least-squares optimization method that iteratively updates the free parameter values to reduce the residual error between simulations and observations (log_10_ viral abundances). When measurement standard errors were available, they were propagated into log_10_ space and used to weight the data points, where less weight was given to data points with more error. If no error bars were given in the data, unweighted least squares was used. Parameters were constrained to biologically realistic ranges, with proportions restricted to [0, 1] and decay rates constrained to positive values.

For the biphasic model (Equation 2), the mixing fraction is initialized using a grid search over 19 values from *p* = 0.05 to 0.95, with the best-fitting solution retained. The estimated initial decay rates, *k*_1_, and *k*_2_, are initialized by fitting lines to early and late portions of the log_10_ time series. These slopes were converted to exponential decay-rate units to obtain *k*_1,0_ and *k*_2,0_, with constraints *k*_1,0_ *> k*_2,0_ *>* 0. For the aggregation model (Equation 5), the intial fraction of single virions *p*_init_ is initialized at the fitted biphasic fraction *p*, which provides an empirically informed starting point based on the biphasic fraction of fast decaying virions. The aggregation rate *f* is initialized by matching the aggregation loss rate per single virion 2*f V*_1_ to the fast biphasic decay scale *k*_1_, giving *f*_init_ = *k*_1_*/*(2*V*_1,init_) with *V*_1,init_ = *p*_init_*V*_0_. Here, *V*_1_ has units of density, *k*_1_ has units of time^*−*1^, and *f* has units of density^*−*1^ time^*−*1^, so 2*f V*_1_ has units of time^*−*1^. For PFU-based fits, the single virion decay rate was initialized as *d*_init_ = (*pk*_1_ + (1 *− p*)*k*_2_)*/p*_init_ (Equation S24), with units of time^*−*1^.

For model comparison between the exponential and biphasic fits, Akaike’s Information Criterion (AIC) was computed directly from the `lmfit` package. As several datasets had few data points (*n*) relative to the number of fitted parameters (*k*) (*n/k <* 40), we used the small-sample correction (AICc) and compared models using ΔAICc, with values *<* 2 treated as indistinguishable, 2 - 4 as moderate support, and *>* 4 as strong support for the lower AICc model [36]. We also report *R*^2^ in log_10_ space as a descriptive measure of goodness of fit. The *R*^2^ values were computed using the `lmfit` package [35], which defines 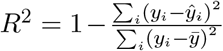. Under this definition, *R*^2^ can be negative when the model predictions are worse than using the mean observed value, so *−∞ < R*^2^ *<* 1.

### Implementation

All analyses were performed in Python 3.1. Nonlinear least-squares fitting and model comparison were implemented using `lmfit` [35], with numerical operations handled by `NumPy` [37]. Figures were generated using `Matplotlib` [38].

## Results

### Biphasic decay is prevalent across viruses of microbes

We identified 17 datasets with sufficiently high temporal resolution and performed AICc model comparisons between exponential decay (Equation 1) and biphasic decay (Equation 2) for each dataset (Figure 1). Using AICc model comparison, we found that biphasic decay was strongly favored in 6 datasets, while exponential decay was strongly favored in 6 datasets (Figure 2). In 2 datasets, Boixereu et al. [39] and Chen et al. [40], AICc model preference was statistically indistinguishable, although the Boixereu et al. dataset (Figure 1f) showed a qualitatively clear biphasic pattern and much better biphasic fit quality 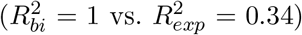 These results show that support for biphasic decay recurs across multiple virus-microbe systems and multiple environments, including soil, gut, plant, river, and oceanic systems. The mixed model support across datasets indicates that exponential decay remains appropriate in some cases, suggesting that extracellular viral decay is both system- and context-dependent. Assuming a single fast exponential decay process across VOM systems may obscure important variation in viral persistence and turnover. Biphasic supported datasets also tended to have more data points than exponential supported datasets, with a median of 9 data points compared to 5, suggesting that more and longer sampling may be important for resolving non-exponential decay dynamics. This finding is apparent in Boixereu et al.[39], where a visually evident biphasic pattern was not strongly supported by AICc as sparse sampling heavily penalized AICc support for the more parameter heavy biphasic model (Refer to *Supplementary Information: Biphasic vs. exponential decay across datasets* for more details on sparse sampling penalization in AICc). Full fitted parameters and statistics are shown in Table S3.

**Figure 1.**
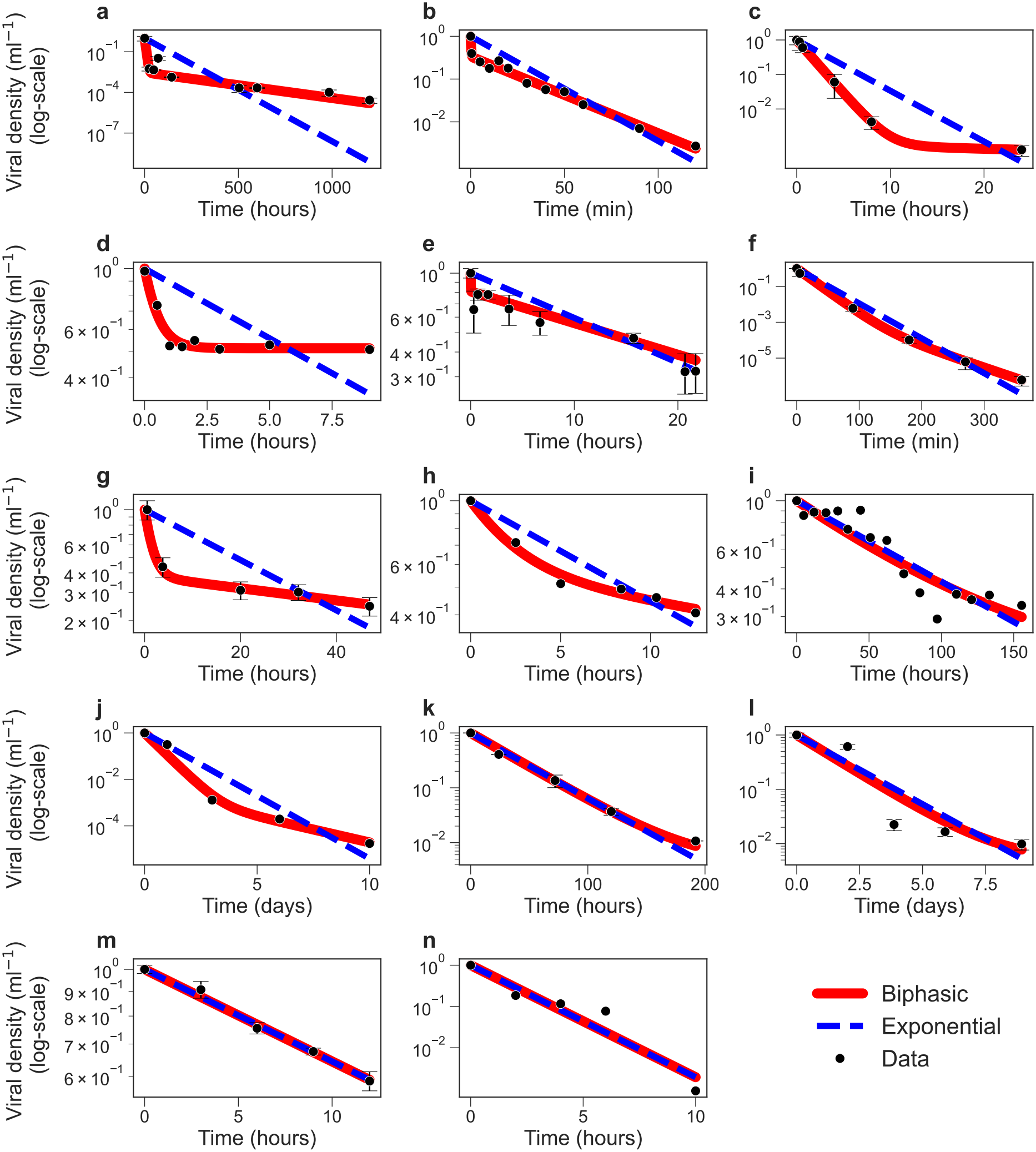
Biphasic decay recurs across multiple virus-microbe systems. Viral decay data from studies identified in the Web of Science search are shown in log-space (points; *±*SEM where available), with best-fit exponential (dashed blue) and biphasic (solid red) models. Fits were performed in log-space after normalizing each dataset by its initial virion density. The environmental habitat of each virus–microbe system is denoted in the top right of each panel.Panels show datasets from: (a) DiPietro et al. 2023 [41]; (b) Shaffer et al. 2024 [42]; (c) French et al. 2023 [43]; (d) Fischer et al. 2002 [44]; (e) Mathias et al. 1995 [45]; (f) Blazanin et al. 2022 [46]; (g) Boixereu et al. 2002 [39]; (h) Chen et al. 2011 [40]; (i) Parada et al. 2007 [47]; (j) Petterson et al. 2001 [29]; (k) Suttle et al. 1992 [48]; (l) Moebus et al. 1992 [49]; (m) Wei et al. 2018 [7]; and (n) Panagiotis et al. 2025 [50]. See Methods: Model Fitting and Table S3 for fitting details and parameter values.

**Figure 2.**
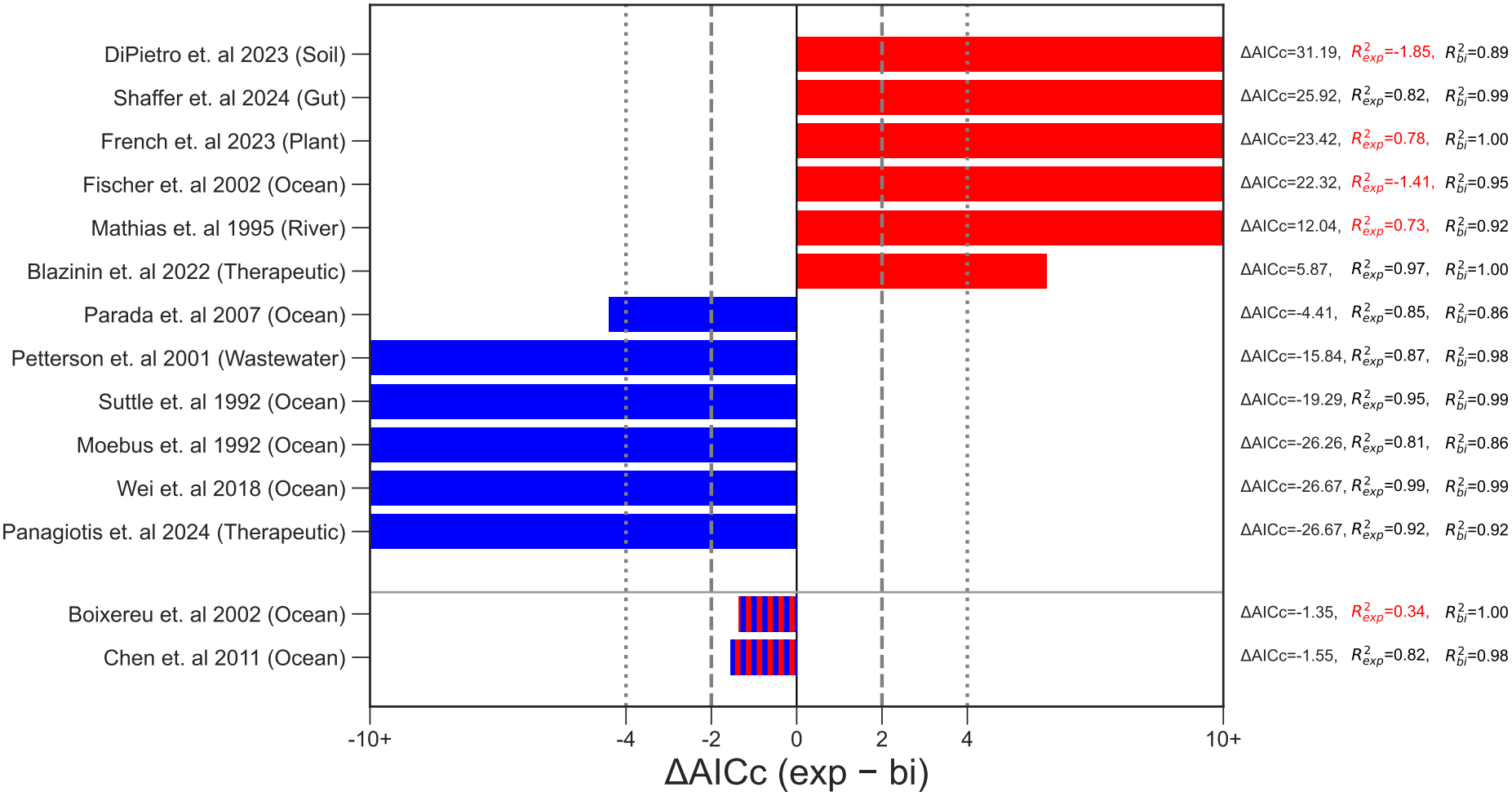
AICc analysis reveals recurring support for biphasic viral decay across datasets. Across all our collected datasets [49, 48, 39, 44, 51, 47, 40, 7, 52, 46, 41, 53, 42, 43, 50, 29, 45], model support is quantified using ΔAICc, defined as AICc_bi_ *−* AICc_bi_. Negative values (blue) indicate support for exponential decay while positive values (red) indicate support for biphasic decay. Vertical dashed lines mark the threshold for indistinguishable support (|ΔAICc| *<* 2) (mixed colors), while dotted lines indicate moderate (2 *≤* |ΔAICc| *≤* 4) and strong (|ΔAICc| *>* 4) support. *R*^2^ values for both the exponential and biphasic fits for each dataset shown on the right axis, where red values fall below the *R*^2^ *≥* 0.8 threshold. Of the 17 datasets analyzed, three datasets (Fischer et al. 2004[51]; Tsai et al. 2022[52]; Pleyer et al. 2024[53]) are excluded from the figure because model fits did not meet the *R*^2^ *≥* 0.8 threshold (either both models or the winning model). The remaining 14 datasets are shown here. All the fitted parameters and statistics are shown in detail in Table S3.

### Extracellular virion aggregation as a proposed generic mechanism that can drive non-exponential decay

The prevalence of slower-than-exponential decay in VOMs raises the question of what biological mechanisms could generate these patterns. In addition to intrinsic heterogeneity, we hypothe-size that generic dynamical processes can produce non-exponential decay across diverse virus-microbe systems. One plausible mechanism is extracellular virion aggregation, which has been linked to non-exponential decay in viruses of humans [23, 22] and observed phenomenologically in viruses of microbes [34, 32, 54]. We therefore developed a mechanistic model of reversible aggregation, in which single virions can form aggregates that are temporarily protected from decay before separating back into single virions:

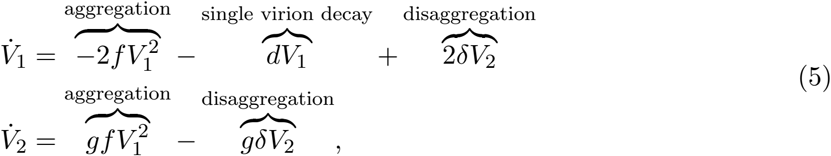

where *V*_2_ aggregates form with aggregation rate *f* from two *V*_1_ virions, and disaggregate back into two *V*_1_ virions at a rate *δ*. We focus on two-virion aggregates rather than higher order multi-virion aggregates; as we show, protected dimers are sufficient to generate slower-than-exponential population-scale decay dynamics without including complex, multi-virion aggregates. Although aggregates could also decay, we assume that the durability of aggregates is far longer than single virions and that direct aggregate decay does not appreciably impact viral density measurements. The parameter *g* accounts for the measurement method, with *g* = 2 for genome based measurements, where both genomes within a dimer aggregate are counted, and *g* = 1 for plaque-forming unit (PFU) based measurements, where a dimer aggregate is counted as a single infectious unit (Figure 3). Note that Equation 5 is the virus particle-associated subset of Equation 4 (given *S* = *I* = 0).

**Figure 3.**
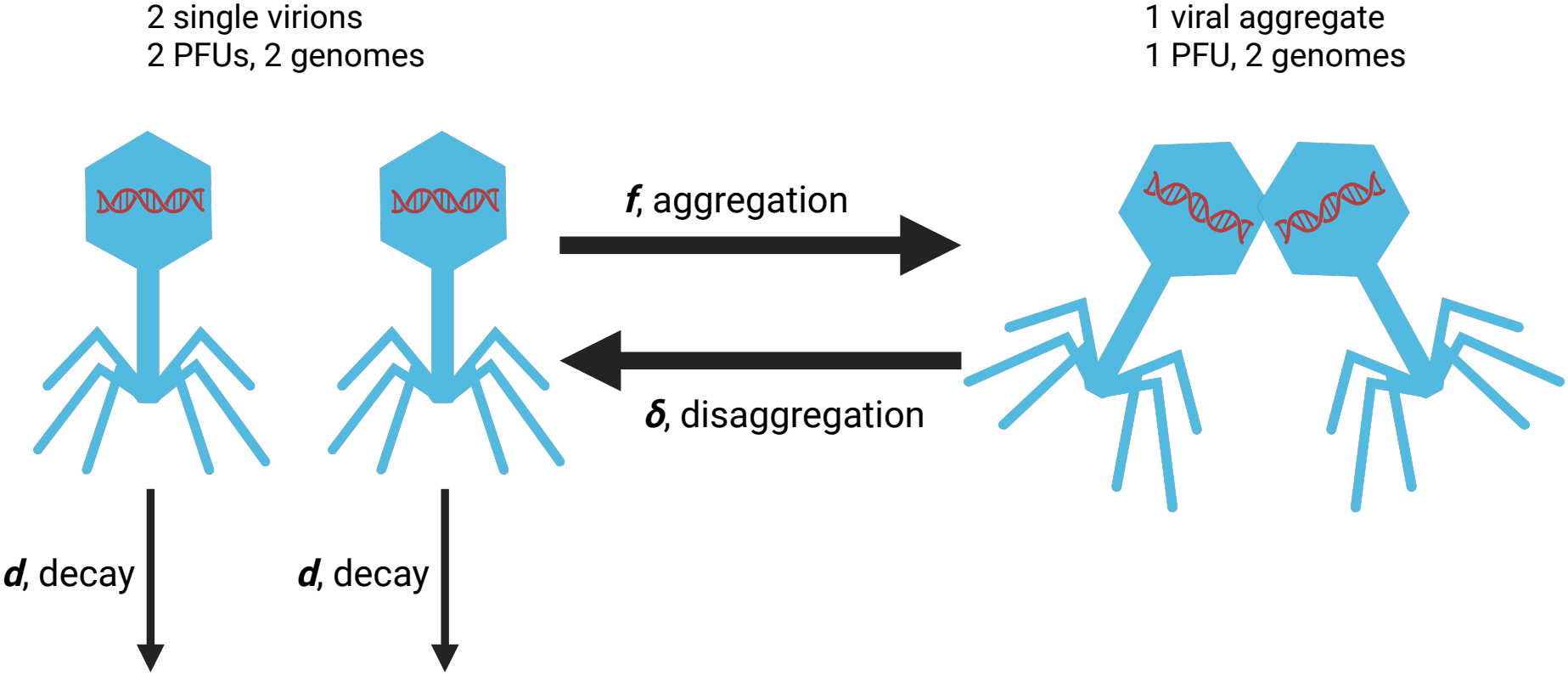
Aggregation and disaggregation alter PFU count but not genome count. Schematic illustration how virion aggregation and disaggregation produce different observed totals dependent on the choice of genome or PFU based measurments. Two single virions correspond to two genomes and two PFUs. Upon aggregation, the two genomes become contained with a single PFU. Subsequent disaggregation restores two PFUs without changing the number of genomes. Therefore PFU count (*g* = 1 in Equation 5) decreases upon aggregation and increases upon disaggregation, while genome count (*g* = 2 in Equation 5) remains conserved.

To relate the mechanistic aggregation model (Equation 5) to the subpopulation-based biphasic formulation (Equation 2), we derived analytical parameter mappings between the model classes. Specifically, the disaggregation rate from the aggregation model *δ* can be directly mapped by the slow decay rate of the biphasic model *k*_2_, allowing part of the aggregation model parameterization to be directly constrained by the corresponding biphasic fit. The remaining aggregation parameters were found to be analytically unidentifiable, where different combinations of parameter values produced the same best-fit (Figure S2). Full details of the analytically mapped and numerically fit parameters are provided in *Supplementary Information: Mathematical derivations of aggregation-to-biphasic mapping* and *Supplementary Information: Biphasic vs aggregation across biphasic-favored datasets*.

Across the datasets found to exhibit biphasic viral decay, the mechanistic aggregation model uniformly reproduce biphasic decay patterns, capturing both the initial rapid decay phase and the subsequent slower persistent phase seen in the corresponding biphasic fits (Figure 4a-f). Boixereu et al [39] was included as well as it shows a visually apparent biphasic pattern despite statistically indistinguishable AICc support (Figure 4g). Together, the repeated observation of biphasic dynamics across datasets and the analytical mapping of a subset of parameters suggest that extracellular aggregation consistently and generically recapitulates biphasic decay.

**Figure 4.**
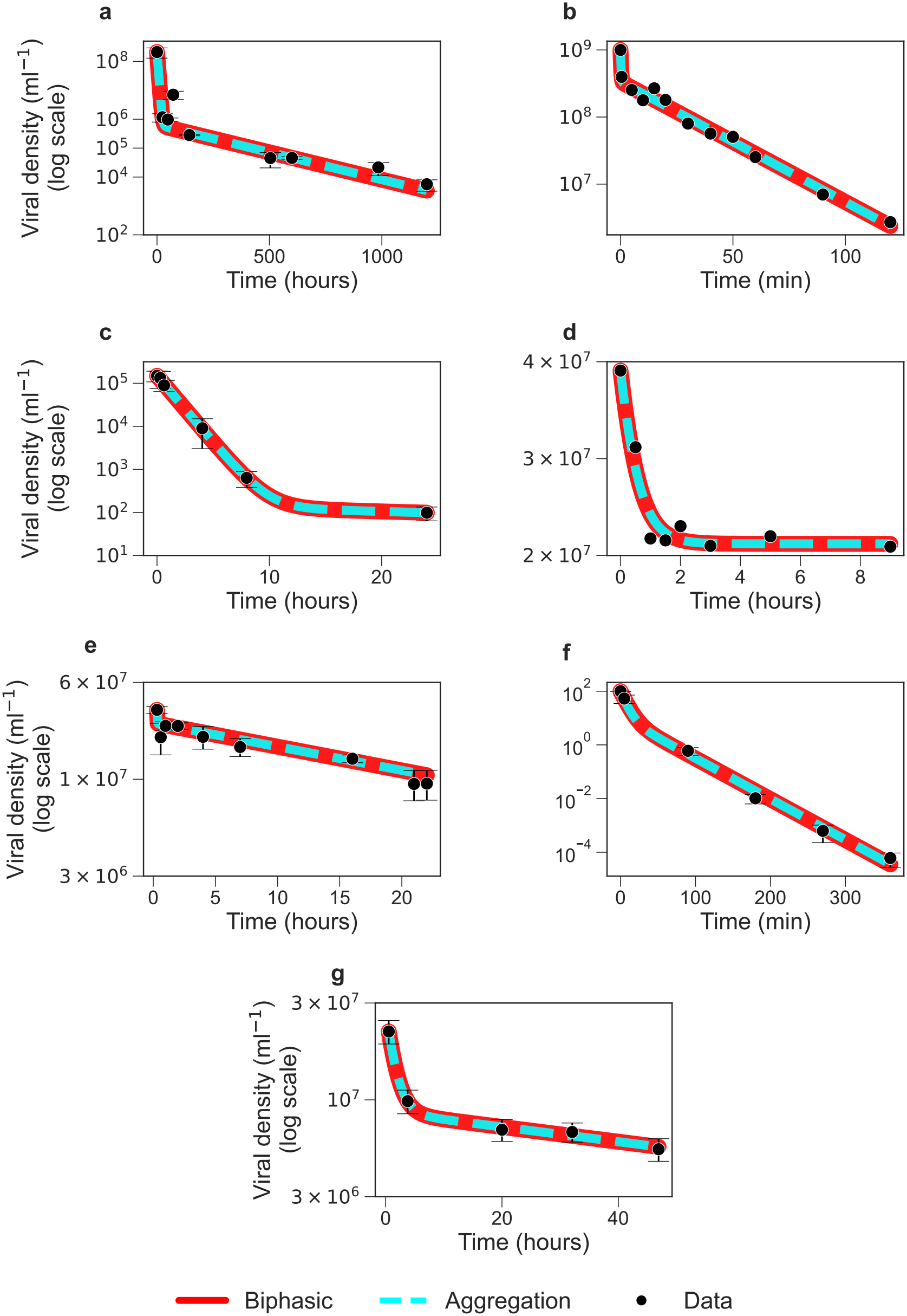
Aggregation model reproduces decay patterns in datasets exhibiting biphasic dynamics. Viral decay data from identified as biphasic are shown in log-space (points; *±*SEM where available), with best-fit biphasic (solid red) and agrgegation (dashed blue) models. For datasets reporting viral genome density ((a) DiPietro et al. 2023 [41], (e) Mathias et al. 1995 [45], and (g) Boixereu et al. 2002 [39]), aggregation model parameters were constrained using the corresponding biphasic fit, with *δ* = *k*_2_*/*2 and *d* = *pk*_1_ + (1 *− p*)*k*_2_, while *f* and *p*_0_ were estimated via nonlinear least squares. For datasets reporting PFU ml^*−*1^ ((b) Shaffer et al. 2024 [42], (c) French et al. 2023 [43], (d) Fischer et al. 2002 [44]), and (f) Blazanin et. al 2022 [46],*δ* = *k*_2_ was fixed from the biphasic fit, and *d, f*, and *p*_0_ were estimated via nonlinear least squares. All fits done in log-space. Refer to Methods: Model Fitting for details on the fitting process and Table S4 for parameter values.

### Extracellular aggregation-induced biphasic decay decouples viral abundance from infection pressure

Rapid aggregation and slow disaggregation would hypothetically allow for virus-host systems to co-exist at higher infection pressures. Aggregation allows virions to enter a persistent state, thereby maintaining high viral abundances through persistence rather than turnover. To evaluate this mechanism, we incorporated reversible aggregation into a nonlinear virus-host population dynamic model that tracks susceptible and infected hosts, free infectious virions, and non-infectious aggregates (Equation 4). To examine this regime, we fixed total viral abundance at 5 *×* 10^8^ virions L^*−*1^ and compared our aggregation model with a similar model with no aggregation (Equation 3) that was calibrated to produce 1% steady state infection as reported across multiple experimental studies of marine virus-host dynamics [19, 18, 55].

We find that independently of variation in adsorption rate *ϕ*, increasing the aggregation rate *f* reduces the steady state fraction of infected hosts, even while total viral abundance remains unchanged (Figure 5a, Equation S47). This is explained by the accompanying reduction in the steady state fraction of free infectious *V*_1_ virions, where as *f* increases, a larger share of the viral population is sequestered into protected non-infectious *V*_2_ aggregates, reducing the infectious subpopulation available to drive new infections while keeping viral abundances high by increasing the number of persistent aggregates (Figure 5b, Equation S69). Thus, high standing viral abundance need not imply high realized infection, but may instead reflect extracellular persistence rather than continual infection-driven viral production and turnover. Likewise, increasing the adsorption efficiency of single virions monotonically increased the steady state infection level. This analysis reveals an identifiability limitation, where similar virus-induced ecological outcomes can arise either from inefficient adsorption by a larger pool of infectious virions or from more efficient adsorption within a smaller infectious subpopulation (Figure 5c, Equation S105). Full details of the analytical steady states and parameter values are provided in *Supplementary Information: Mathematical derivations of virus-host model steady states* and Table S5 respectively.

**Figure 5.**
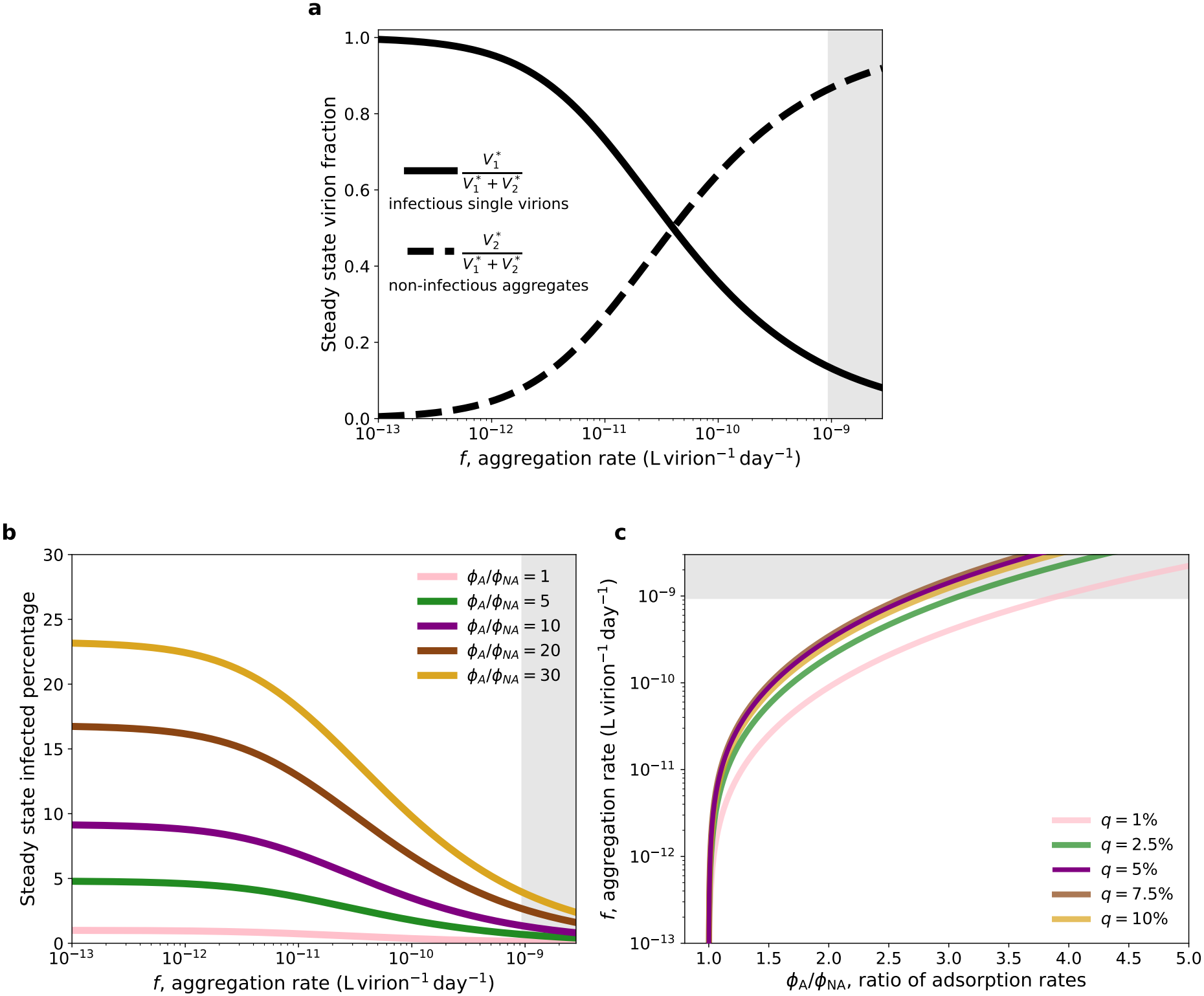
Sufficiently high rates of extracellular aggregation suppress infection while maintaining high viral abundance. Total steady state viral abundance is fixed at 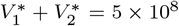 virions L^*−*1^ [18, 56]. Gray region denotes aggregation rates exceeding the Stokes-Einstein biophysical limit for virus-virus contact rates. **(a)** Steady state fractions of infectious single virions, 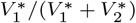 (Equation S47), and non-infectious aggregates, 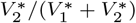 (Equation S48), as functions of the aggregation rate *f*. Increasing aggregation progressively shifts the viral population from infectious single virions to non-infectious aggregates. This result is independent of variation in *ϕ* **(b)** Steady state infected-host percentage as a function of *f* for increasing adsorption-rate ratios *ϕ*_A_*/ϕ*_NA_ (Equation S69), where *ϕ*_NA_ is fixed to be = 9.61 *×* 10^*−*11^ L virion^*−*1^ day^*−*1^ to yield *q* = 1% steady state infection in the no aggregation model, and *ϕ*_A_ is the adsorption rate in the aggregation model. Greater adsorption in the aggregation model partially compensates for the loss of infectious single virions, whereas increasing aggregation suppresses infection. **(c)** Combinations of *ϕ*_A_*/ϕ*_NA_ and *f* that preserve the same total steady state viral abundance for fixed steady state infected host percentages *q* (Equation S105). Higher aggregation therefore requires compensatory increases in adsorption efficiency to maintain a given infection prevalence, illustrating parameter non-identifiability between the aggregation rate and the adsorption rate. Full details of the virus-host models are provided in *Methods: Virus-Host Models*, parameter values are provided in Table S5, and steady states are provided in *Supplementary Information: Mathematical derivations of virus-host model steady states*

## Discussion

Conventional virus-microbe models typically assume rapid exponential decay of extracellular virions. Under this assumption, high viral abundances are often interpreted as evidence for elevated rates of host infection and mortality to maintain constant virion turnover. Here, we reassessed this assumption by showing that, across 17 published viral decay datasets, biphasic decay, reflecting fast and slow decaying viral subpopulations, is statistically supported approximately as frequently as a single exponential decay model. We further show that extracellular virion aggregation can reproduce biphasic decay, with aggregation model parameters directly mappable onto the parameters of the biphasic decay model. This correspondence suggests that biphasic decay need not arise solely from fixed heterogeneity among virions, but can also emerge dynamically from ecological and/or biophysical processes. Overall, these results challenge conventional assumptions that uniformly rapid exponential decay and high viral turnover are required to maintain observed viral abundances. Instead, if a fraction of extracellular virions persist durably, then lower rates of host infection, killing, and virion turnover may be sufficient to sustain the observed levels of standing viral abundances.

These findings address a longstanding tension in virus-microbe ecology that physical encounter-rate arguments predict frequent virus-host contacts, yet realized infection rates are often much lower than expected, even where viral abundances remain high [18, 19, 55, 54]. Standard models with rapid exponential decay typically reconcile this pattern by inferring low bulk adsorption efficiency across the entire viral population. To examine whether implementing slower-than-exponential decay can offer an alternative explanation, we incorporated aggregation into a nonlinear virus-host model and compared its steady state dynamics with an equivalent model lacking aggregation. We found that aggregation shifted virions from the free infectious pool into a persistent non-infectious pool, reducing infection pressure and thereby the fraction of infected hosts while total viral abundance was held constant. Furthermore, we find that higher adsorption efficiency among the remaining free virions could compensate for their reduced abundance, allowing similar infection levels to arise from different combinations of adsorption efficiency and infectious pool size. Together, our results suggest that frequent physical encounters and high viral abundance can coexist with weak realized infection because all virions adsorb inefficiently or, alternatively, because a substantial fraction of the standing viral pool is persistent and non-infectious. These results suggest the need for direct measurements of viral aggregates and, potentially, a reassessment of adsorption assays that neglect heterogeneity [57, 58, 59, 60].

Our study has several caveats. We focused primarily on fast and slow decaying viral sub-populations, and on a single dynamical mechanism in aggregation as drivers of biphasic decay. Several alternative mechanisms could generate apparent fast and slow decaying subpopulations. For example, extracellular vesicles have been shown to increase viral persistence in the environment [61, 62], while vesicle enclosure may also reduce receptor binding and host infection [61]. In addition, environmental and experimental conditions, such as temperature, light exposure, humidity, and adsorption to particles or surfaces, can strongly affect virion stability and life history [7, 60, 63, 64]. Moreover, infectivity may be restored through light-dependent photoreactivation or host-mediated dark repair pathways [65, 66]. These different mechanisms and environmental factors may have distinct implications for virus-host population dynamics, that are not distinguishable by decay time series alone. Furthermore, the present study only considers two-virion aggregates even if multi-virion aggregates are possible [34, 22, 32]. Nevertheless, a model with protected two-virion aggregates is sufficient to generate slower-than-exponential decay dynamics at the population level; we expect that including multi-virion aggregates would enhance this effect [9]. In addition, our aggregation models assume that genome counts are conserved during aggregation and disaggregation, which holds only if these processes do not disrupt viral genomes. Similarly, our interpretation of PFU dynamics assumes that aggregation removes one measurable PFU and disaggregation restores one measurable PFU only if these processes do not independently alter infectiousness. Together, these assumptions highlight the need for future experimental work to determine how viral (dis)aggregation affects virion persistence and their interactions with host cells [67].

Altogether these findings show that slower-than-exponential extracellular decay is commonplace within VOMs and can decouple viral standing abundance from viral turnover. When a fraction of virions persists longer than expected under a single exponential decay model, then lower levels of viral-induced host lysis may be sufficient to sustain abundant viral populations. As a result, extracellular virion persistence may be an underappreciated ecological mechanism shaping virus-microbe coexistence and the interpretation of high viral abundances in nature.

## Supporting information

Suppelementary Information

## Author Contributions

A.A, E.W., and J.S.W conceptualized the study. A.A, and E.W. performed the decay data collection. A.A and P.F. performed the analytical mathematical analysis. A.A. conducted the simulations. A.A. conducted the statistical analysis and visualization. A.A, P.F., and J.S.W prepared the original draft of the manuscript. All authors contributed to the writing and approval of the final version of the manuscript.

## Conflicts of interest

None declared

## Funding

This project was supported by Simons Foundations grants SFI-LS-PROJECT-00011964 and MPS-SIP-00930382 awarded to J.S.W. J.S.W. is an investigator at the University of Maryland-Institute for Health Computing, which is supported by funding from Montgomery County, Maryland, and The University of Maryland Strategic Partnership: MPowering the State, a formal collaboration between the University of Maryland, College Park, and the University of Maryland, Baltimore.

## Acknowledgements

We sincerely thank Emily Bruns, Steve Wilheim, Claudia Igler, Drew Thornley, and Raunak Dey for their comments and feedback on revisions of the manuscript. We also thank Kejia Zhang for reviewing the code and suggesting improvements.

## Data accessibility statement

The viral decay datasets analyzed in this study originate from the 17 published papers listed in Table S3. The datasets and code used for data analysis and figure generation are also openly accessible at https://github.com/aranilah/Slower-than-exponential-viral-decay.

## Notes

https://github.com/aranilah/Slower-than-exponential-viral-decay

### Competing Interest Statement

The authors have declared no competing interest.

### Summary of Updates

Corrected typo in Figure 2 caption

