## Supplementary material for "Slower-than-exponential viral decay is prevalent and can reshape virus-microbe dynamics": Suppelementary Information

### Supplementary Information

#### Supplementary Tables

| Symbol | Units | Definition |
|---|---|---|
| $V(t)$ | abundance | Viral abundance at time $t$ |
| $V_0$ | abundance | Viral abundance at the start of the experiment, $t = 0$ (Normalized in most cases to $V_0 = 1$) |
| $t$ | time | Time since the start of the decay experiment |
| $k$ | $\text{time}^{-1}$ | Decay rate constant in the monophasic exponential model |
| $p$ | dimensionless | Proportion of the viral population decaying at rate $k_1$ |
| $1 - p$ | dimensionless | Proportion of the viral population decaying at rate $k_2$ |
| $k_1$ | $\text{time}^{-1}$ | Exponential decay rate of the faster decaying viral subpopulation |
| $k_2$ | $\text{time}^{-1}$ | Exponential decay rate of the slower decaying viral subpopulation |

**Table S1.** Parameters and state variables in the exponential (Equation 1) and biphasic (Equation 2) decay models.

| Symbol | Units | Definition |
|---|---|---|
| $V_1$ | virions/L | Density of single virions |
| $V_2$ | virions/L | Density of aggregates |
| $V_{\text{tot}}$ | virions/L | Total viral density, $V_1 + V_2$ |
| $f$ | $\text{abundance}^{-1}$ $\text{time}^{-1}$ | Rate at which single virions collide and form aggregates |
| $\delta$ | $\text{time}^{-1}$ | Rate at which aggregates dissociate back into single virions |
| $d$ | $\text{time}^{-1}$ | Exponential decay rate of single, unaggregated virions |
| $t$ | time | Time since the start of the decay experiment |
| $g$ | unitless | viral genome measurement parameter |

**Table S2.** Parameters and state variables in the host-free aggregation model (Equation 5).

| Paper | Native habitat | $\Delta$AICc($\text{AICc}_{\text{exp}} - \text{AICc}_{\text{bi}}$) | $R^2_{\text{exp}}$ | $R^2_{\text{bi}}$ | $k\,(time^{-1})$ | $p$ | $k_1\,(time^{-1})$ | $k_2\,(time^{-1})$ |
|---|---|---|---|---|---|---|---|---|
| DiPietro et al. 2023(DiPietro et al., 2023) | Soil | 31.188 | -1.849 | 0.894 | $0.018 \pm 0.004$ | $0.997 \pm 0.001$ | $0.240 \pm 0.107$ | $0.004 \pm 0.001$ |
| Shaffer et al. 2024(Shaffer et al., 2024) | Gut | 25.915 | 0.818 | 0.988 | $0.056 \pm 0.004$ | $0.664 \pm 0.035$ | $4.574 \pm 2.613$ | $0.041 \pm 0.002$ |
| French et al. 2023(French et al., 2023) | Human | 23.422 | 0.782 | 1.000 | $0.339 \pm 0.049$ | $0.999 \pm 0.002$ | $0.712 \pm 0.325$ | $0.016 \pm 0.070$ |
| Fischer et al. 2002(Fischer and Velimirov, 2002) | Ocean | 22.322 | -1.412 | 0.954 | $0.117 \pm NA$ | $0.488 \pm NA$ | $2.115 \pm NA$ | $3.375^{-14} \pm NA$ |
| Mathias et al. 1995(Mathias et al., 1995) | Wastewater/Feces/Soil | 12.045 | 0.725 | 0.916 | $0.052 \pm 0.006$ | $0.192 \pm 0.024$ | $71.719 \pm NA$ | $0.037 \pm 0.003$ |
| Blazanin et al. 2022(Blazanin et al., 2022) | Human | 5.867 | 0.966 | 0.995 | $0.045 \pm 0.004$ | $0.990 \pm 0.008$ | $0.056 \pm 0.003$ | $0.027 \pm 0.003$ |
| Parada et al. 2007(Parada et al., 2007) | Ocean | -4.407 | 0.848 | 0.860 | $0.008 \pm 0.001$ | $0.831 \pm NA$ | $0.012 \pm NA$ | $10^{-15} \pm NA$ |
| Petterson et al. 2001(Petterson et al., 2001) | Wastewater | -15.842 | 0.867 | 0.985 | $1.246 \pm 0.140$ | $0.997 \pm 0.008$ | $2.179 \pm 0.427$ | $0.511 \pm 0.291$ |
| Suttle et al. 1992(Suttle and Chen, 1992) | Ocean | -19.288 | 0.954 | 0.993 | $0.027 \pm 0.000$ | $0.995 \pm 0.016$ | $0.028 \pm 0.001$ | $10^{-12} \pm NA$ |
| Moebus et al. 1992(Moebus, 1992) | Ocean | -26.260 | 0.810 | 0.861 | $0.586 \pm 0.090$ | $0.995 \pm 0.232$ | $0.657 \pm 0.531$ | $10^{-12} \pm NA$ |
| Wei et al. 2018(Wei et al., 2018) | Ocean | -26.667 | 0.992 | 0.992 | $0.044 \pm 0.001$ | $1.00 \pm NA$ | $0.044 \pm NA$ | $0.067 \pm NA$ |
| Panagiotis et al. 2024(Zagaliotis et al., 2025) | Human | -26.667 | 0.916 | 0.916 | $0.625 \pm 0.061$ | $10^{-12} \pm NA$ | $22.036 \pm NA$ | $0.625 \pm NA$ |
| Boixereu et al. 2002(Guixa-Boixereu et al., 2002) | Ocean | -1.345 | 0.345 | 0.997 | $0.036 \pm 0.007$ | $0.619 \pm 0.003$ | $0.688 \pm 0.158$ | $0.009 \pm 0.003$ |
| Chen et al. 2011(Chen et al., 2011) | Ocean | -1.545 | 0.817 | 0.981 | $0.081 \pm 0.007$ | $0.436 \pm 0.205$ | $0.419 \pm 0.279$ | $0.024 \pm 0.030$ |
| Fischer et al. 2004(Fischer et al., 2004) | Ocean | -2.583 | -0.305 | 0.835 | $0.102 \pm 0.028$ | $0.305 \pm 0.150$ | $1.63 \pm 2.193$ | $10^{-12} \pm NA$ |
| Tsai et al. 2022(Tsai et al., 2022) | River | -4.629 | -0.200 | 0.330 | $0.027 \pm 0.004$ | $0.357 \pm 1.471$ | $0.123 \pm 0.468$ | $10^{-3} \pm NA$ |
| Pleyer et al. 2024(Pleyer et al., 2024) | River | -6.303 | 0.733 | 0.733 | $0.039 \pm 0.004$ | $0.746 \pm NA$ | $0.039 \pm NA$ | $0.039 \pm NA$ |

**Table S3.** Fitted parameters and statistics used in Figure 1 and Figure 2. The datasets from Fischer et al. 2004 (Fischer et al., 2004), Tsai et al. (Tsai et al., 2022), and Pleyer et al. (Pleyer et al., 2024) are shown at the end because they were excluded from the main text but shown in Figure S1 as the $R^2$ values of the winning fit, or of both fits, fell below the 0.8 threshold. All analysis was performed on normalized datasets. Native habitat indicates the environmental or host-associated context from which each virus-microbe system was drawn. $R^2$ values are reported for exponential and biphasic fits. $NA$ indicates that parameter uncertainty could not be reliably estimated because the fitted parameter was at or near a boundary or was poorly identifiable.

| Paper | $p$ | $k_1\,(time^{-1})$ | $k_2\,(time^{-1})$ | $p_0$ | $f(concentration^{-1} \times time^{-1})$ | $\delta(time^{-1})$ | $d(time^{-1})$ |
|---|---|---|---|---|---|---|---|
| DiPietro et al. 2023(DiPietro et al., 2023) | $0.997 \pm 0.001$ | $0.240 \pm 0.107$ | $0.004 \pm 0.001$ | 0.999 | $2.840 \times 10^{-12}$ | $k_2/2$ | $\frac{pk_1+(1-p)k_2}{p_0}$ |
| French et al. 2023(French et al., 2023) | $1.000 \pm 0.002$ | $0.712 \pm 0.033$ | $0.016 \pm 0.070$ | 1.00 | $8.778 \times 10^{-9}$ | $k_2$ | 0.712 |
| Shaffer et al. 2024(Shaffer et al., 2024) | $0.664 \pm 0.035$ | $4.574 \pm 2.614$ | $0.041 \pm 0.002$ | 0.667 | $1.634 \times 10^{-13}$ | $k_2$ | 4.574 |
| Fischer et al. 2002(Fischer and Velimirov, 2002) | $0.476 \pm 0.034$ | $2.053 \pm 0.525$ | $3.891 \times 10^{-9}$ | 0.476 | $5.773 \times 10^{-15}$ | $k_2$ | 2.053 |
| Mathias et al. 1995(Mathias et al., 1995) | $0.192 \pm 0.034$ | $73.560 \pm 0.000$ | $0.037 \pm 0.003$ | 0.346 | $2.728 \times 10^{-6}$ | $k_2/2$ | $\frac{pk_1+(1-p)k_2}{p_0}$ |
| Blazanin et al. 2022(Blazanin et al., 2022) | $0.898 \pm 0.006$ | $0.138 \pm 0.103$ | $0.035 \pm 0.003$ | 1 | $1.81 \times 10^{-4}$ | $k_2$ | 0.144 |
| Boixereu et al. 2002(Guixa-Boixereu et al., 2002) | $0.619 \pm 0.033$ | $0.688 \pm 0.158$ | $0.009 \pm 0.004$ | 0.624 | $7.075 \times 10^{-12}$ | $k_2/2$ | $\frac{pk_1+(1-p)k_2}{p_0}$ |

**Table S4.** Comparison of biphasic and aggregation model parameters across datasets exhibiting strong support for biphasic decay. Datasets are ordered by descending ΔAICc. While all viral data was normalized in Table S3, all fits shown here are performed on non-normalized data. As a result, parameter estimates may sometimes differ quantitatively from the corresponding values reported for normalized fits. In Fischer et. al 2002, the fitted slow decay rate $k_2$ is effectively zero and poorly constrained by the data, thereby covariance-based standard error is not meaningful. No errors are reported for the aggregation parameters as they are either analytically mapped or are non-identifiable Figure S2.

| Symbol | Units | Definition | Figure 5a,b | Figure 5c |
|---|---|---|---|---|
| $S^*$ | cells $L^{-1}$ | steady state density of susceptible host cells | Analytically determined | Analytically determined |
| $I^*$ | cells $L^{-1}$ | steady state density of infected host cells | Varies with $f$ and $\phi_A/\phi_{NA}$ | Fixed through specified $q$ |
| $V^*/V_{target}$ | virions $L^{-1}$ | steady state total viral abundance | $5 \times 10^8$ | $5 \times 10^8$ |
| $V_1^*$ | virions $L^{-1}$ | steady state density of free infectious virions in the aggregation model | Varies with $f$; but follows $V_1^* + V_2^* = 5 \times 10^8$ | Varies with $\phi_A/\phi_{NA}$; but follows $V_1^* + V_2^* = 5 \times 10^8$ |
| $V_2^*$ | virions $L^{-1}$ | steady state density of protected non-infectious aggregates | Varies with $f$; but follows $V_1^* + V_2^* = 5 \times 10^8$ | Varies with $\phi_A/\phi_{NA}$; but follows $V_1^* + V_2^* = 5 \times 10^8$ |
| $r$ | day$^{-1}$ | Maximum host growth rate | Not required | 1.0 |
| $K$ | cells $L^{-1}$ | Carrying capacity for total host density, $S + I$ | Not required | $7.76 \times 10^8$ |
| $\phi_{NA}$ | L virion$^{-1}$ day$^{-1}$ | Adsorption rate in the no-aggregation model | $9.61 \times 10^{-11}$, yielding $q_0 = 0.01$ | Fixed for each $q$ at Equation S75 |
| $\phi_A$ | L virion$^{-1}$ day$^{-1}$ | Adsorption rate of free infectious $V_1$ virions in the aggregation model | Varied to give $\phi_A/\phi_{NA} = 1, 5, 10, 20, 30$ | Varied continuously along each fixed $q$ curve |
| $\omega$ | day$^{-1}$ | Basal loss rate of susceptible and infected host cells | 0.257 | 0.257 |
| $\lambda$ | day$^{-1}$ | Rate at which infected host cells lyse | 4.5 | 4.5 |
| $\beta$ | virions cell$^{-1}$ | Average number of virions released per lysed infected cell | Not required | 24 |
| $d_{NA}$ | day$^{-1}$ | Extracellular virion decay rate in the no-aggregation model | Not required | Equation S82 |
| $d_A$ | day$^{-1}$ | Extracellular virion decay rate of free infectious $V_1$ virions in the aggregation model | 1 | Equation S100 |
| $f$ | L virion$^{-1}$ day$^{-1}$ | Aggregation rate of free infectious virions | Varied from $10^{-13}$ to the biophysical limit, $\sim 9 \times 10^{-10}$ | Varied from $10^{-13}$ to the biophysical limit, $\sim 9 \times 10^{-10}$ |
| $n$ | dimensionless | Scaling factor reducing the disaggregation rate of aggregates | 100 | 100 |
| $q_0$ | dimensionless | Reference steady state infected fraction, $I^*/(S^* + I^*)$, at $f = 0$ and $\phi_A/\phi_{NA} = 1$ | 0.01 | — |
| $q$ | dimensionless | steady state infected fraction, $I^*/(S^* + I^*)$ | Varies with $f$ and $\phi_A/\phi_{NA}$ | Fixed at $0.01,\ 0.025,\ 0.05,\ 0.075,\ 0.10$ |
| $\delta$ | day$^{-1}$ | Disaggregation rate | $\delta = d_A/n$ | $\delta = d_A/n$ |

**Table S5.** Parameters and steady state variables used in the no-aggregation Equation 3 and aggregation Equation 4 virus-host models for Figure 5. Parameters with explicit values were chosen to be broadly consistent with ranges reported across marine virus-host studies (Mruwat et al., 2021; Beckett et al., 2024; Demory et al., 2020)

### Supplementary Figures

#### Biphasic vs. exponential additional datasets

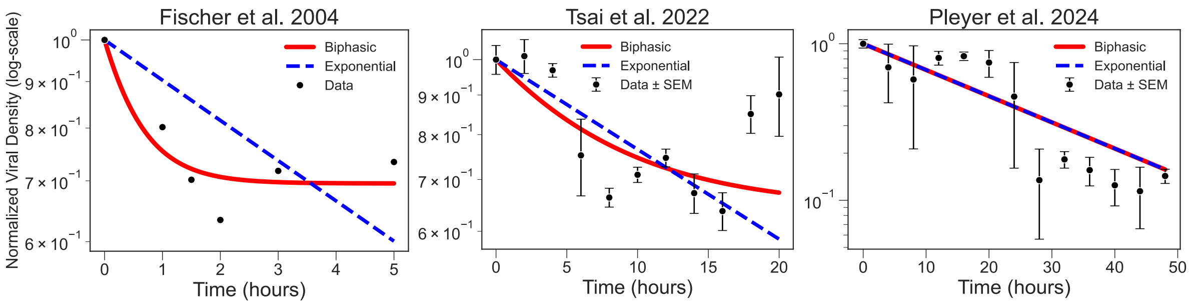


**Figure S1.** Excluded datasets from AICc analysis in main text (Fischer et al., 2004; Tsai et al., 2022). Normalized viral decay data in log-space (points; ±SEM where available) are shown with best-fit exponential (red dashed) and biphasic (solid gray) models. Optimized parameters for each fit are reported in the legend of each panel. All fits done in log-space, and facilitate visual and parameter comparisons across studies, we normalized each dataset by its initial virion density and fixed $V_0 = 1$ in both models. Refer to Methods: Model Fitting for details on the fitting process and Table S3 for parameter values.

#### Parameter identifiability

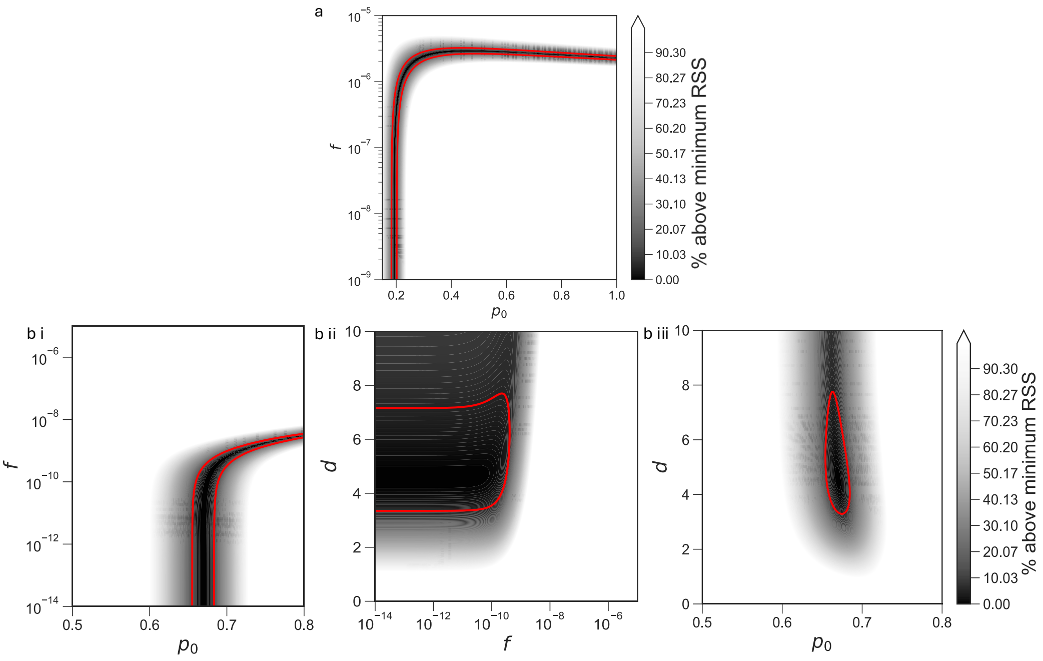


**Figure S2. Parameter non-identifiability analysis in aggregation model fits yields different parameter combinations producing nearly indistinguishable fits.** Heatmap shading shows the percent increase in RSS relative to the minimum RSS, and red contours mark parameter combinations within 5% of the minimum. a) Mathias et al.(Mathias et al., 1995), using genome-based density measurements, exhibits a broad near-optimal region in $(p_0, f)$. b) Shaffer et al. (Shaffer et al., 2024), using PFU-based density measurements, exhibits broad near-optimal regions across pairwise projections of $(p_0, f, d)$. b i) shows $p_0$ vs. $f$ with $d$ fixed, b ii) shows $f$ vs. $d$ with $p_0$ fixed, and b iii) shows $p_0$ vs. $d$ with $f$ fixed; where all the fixed parameters are set to their best-fit value from Table S4.

### Supplementary Text

#### Mathematical derivations of aggregation-to-biphasic mapping

To assess whether aggregation can be analytically linked to biphasic decay, we first modify the biphasic model given by Equation 2 to now have a defined initial virion concentration $V_0$ :

$$V_{\rm bi}(t) = V_0\big[p\, e^{-k_1 t} + (1-p)\, e^{-k_2 t}\big], \qquad k_1 > k_2 > 0. \tag{S1}$$

We set both the genome and PFU formulations of the aggregation model (Equation 5) to have initial conditions $V_1(0) = p_0 V_0, V_2(0) = (1-p_0)V_0$ where $p_0$ is the initial fraction of single virions. We aim to find aggregation parameters $(f, d, \delta, p_0)$ as functions of the biphasic parameters $(V_0, p, k_1, k_2)$ from Equation S1.

We compare the biphasic and aggregation models using the instantaneous per-capita decay rate

$$r(t) = -\frac{\dot{V}_{\rm tot}(t)}{V_{\rm tot}(t)}. \tag{S2}$$

For the biphasic model, we find that the instantaneous decay rate is:

$$r_{\mathrm{bi}}(t)=\frac{pk_1e^{-k_1t}+(1-p)k_2e^{-k_2t}}{pe^{-k_1t}+(1-p)e^{-k_2t}}. \tag{S3}$$

This expression reveals two limiting regimes that constrain any aggregation model reproducing biphasic decay:

- Early-time ($t \approx 0$) limit:

$$r_{\mathrm{bi}}(0)=pk_1+(1-p)k_2, \tag{S4}$$

- Late-time ($t \to \infty$) limit:

$$\lim_{t\to\infty} r_{\mathrm{bi}}(t)=k_2, \tag{S5}$$

Consequently, for the aggregation models to generate biphasic-like dynamics in the total population $V_{\mathrm{tot}}$, they must reproduce instantaneous decay rates that match these early- and late-time limits. Enforcing these two constraints therefore restricts feasible combinations of $(f, d, \delta, p_0)$.

We first consider the genome variant ($g = 2$) of the aggregation model (Equation 5), where summing the differential equations for singles and aggregates yields

$$\dot{V}_{tot}=\dot{V}_1+\dot{V}_2=-dV_1, \tag{S6}$$

$$r_{\mathrm{agg}}(t)=-\frac{\dot{V}_{tot}(t)}{V_{tot}(t)}=\frac{dV_1(t)}{V_{tot}(t)}. \tag{S7}$$

For large $t$, since $V_1$ is small, the quadratic term $2fV_1^2$ can be neglected:

$$\begin{aligned}\dot{V}_2 &\approx -2\delta V_2,\\ \dot{V}_1 &\approx -dV_1+2\delta V_2.\end{aligned} \tag{S8}$$

$$V_2(t)=Ae^{-2\delta t} \tag{S9}$$

$$\dot{V}_1+dV_1=2\delta Ae^{-2\delta t}, \tag{S10}$$

$$V_1(t)=\frac{2\delta A}{d-2\delta}e^{-2\delta t}+Ce^{-dt}, \tag{S11}$$

$$\therefore V_{tot}(t)=V_1(t)+V_2(t)=\left(\frac{2\delta A}{d-2\delta}+A\right)e^{-2\delta t}+Ce^{-dt}=\alpha e^{-2\delta t}+\beta e^{-dt}, \tag{S12}$$

where $\alpha, \beta$ are constants. Assuming biologically that decay of singles is faster than effective loss from aggregates, we take

$$d > 2\delta. \tag{S13}$$

$$\therefore V_{tot}(t) = e^{-2\delta t}\left[\alpha + \beta e^{-(d-2\delta)t}\right], \tag{S14}$$

$$\dot{V}_{tot}(t) = e^{-2\delta t}\left[-2\delta\alpha - d\beta e^{-(d-2\delta)t}\right], \tag{S15}$$

$$r_{\text{agg}}(t) = -\frac{\dot{V}_{tot}(t)}{V_{tot}(t)} = \frac{2\delta\alpha + d\beta e^{-(d-2\delta)t}}{\alpha + \beta e^{-(d-2\delta)t}}. \tag{S16}$$

As $t \to \infty$, $e^{-(d-2\delta)t} \to 0$ and

$$\lim_{t\to\infty} r_{\text{agg}}(t) = 2\delta. \tag{S17}$$

Matching to the biphasic late time limit (Equation S5),

$$\lim_{t\to\infty} r_{\text{agg}}(t) = \lim_{t\to\infty} r_{\text{bi}}(t) = k_2, \tag{S18}$$

$$\boxed{\delta = \frac{k_2}{2}.} \tag{S19}$$

At $t = 0$ we have

$$V_{tot}(0) = V_0, \qquad V_1(0) = p_0 V_0, \tag{S20}$$

$$\therefore \dot{V}_{tot}(0) = -dV_1(0) = -dp_0 V_0. \tag{S21}$$

The instantaneous decay rate at $t = 0$ is

$$r_{\text{agg}}(0) = -\frac{\dot{V}_{tot}(0)}{V_{tot}(0)} = \frac{dp_0 V_0}{V_0} = dp_0. \tag{S22}$$

Matching to the early time decay rates of the biphasic model gives(Equation S4):

$$r_{\text{agg}}(0) = r_{\text{bi}}(0) \quad \Rightarrow \quad dp_0 = pk_1 + (1-p)k_2, \tag{S23}$$

$$\boxed{d = \frac{pk_1 + (1-p)k_2}{p_0}.} \tag{S24}$$

Therefore for the genome model, we only have an exact analytical mapping for $\delta$ and semi-analytical mapping for $d$ (function of $p_0$), and numerically fit $f$, and $p_0$.

We next consider the PFU variant ($g = 1$) of the aggregation model (Equation 5), where summing the differential equations for singles and aggregates this time yields:

$$\dot{V}_{tot} = \dot{V}_1 + \dot{V}_2 = -fV_1^2 - dV_1 + \delta V_2. \tag{S25}$$

For large $t$, we once again drop the quadratic term $fV_1^2$ as $V_1$ is assumed to be small, giving

$$\begin{aligned} \dot{V}_2 &\approx -\delta V_2, \\ \dot{V}_1 &\approx -dV_1 + 2\delta V_2, \end{aligned} \tag{S26}$$

where the solution of this system is

$$V_2(t) = Ae^{-\delta t}, \tag{S27}$$

$$V_1(t) = \frac{2\delta A}{d-\delta}e^{-\delta t} + Ce^{-dt}, \tag{S28}$$

$$\therefore V_{tot}(t) = \alpha e^{-\delta t} + \beta e^{-dt}. \tag{S29}$$

Assuming $d > \delta$,

$$V_{tot}(t) = e^{-\delta t}\left[\alpha + \beta e^{-(d-\delta)t}\right], \tag{S30}$$

$$\dot{V}_{tot}(t) = e^{-\delta t}\left[-\delta\alpha - d\beta e^{-(d-\delta)t}\right], \tag{S31}$$

$$\therefore r_{\text{agg}}(t) = -\frac{\dot{V}_{tot}(t)}{V_{tot}(t)} = \frac{\delta\alpha + d\beta e^{-(d-\delta)t}}{\alpha + \beta e^{-(d-\delta)t}}. \tag{S32}$$

As $t \to \infty$, $e^{-(d-\delta)t} \to 0$

$$\therefore \lim_{t\to\infty} r_{\text{agg}}(t) = \delta. \tag{S33}$$

Matching to the biphasic late time limit (Equation S5):

$$\boxed{\delta = k_2.} \tag{S34}$$

At $t = 0$ we have

$$V_1(0) = p_0 V_0, \qquad V_2(0) = (1 - p_0) V_0. \tag{S35}$$

The time derivative of the total is

$$\dot{V}_{tot}(0) = -f p_0^2 V_0^2 - d p_0 V_0 + \delta(1 - p_0) V_0, \tag{S36}$$

$$r_{\mathrm{agg}}(0) = -\frac{\dot{V}_{tot}(0)}{V_{tot}(0)} = f p_0^2 V_0 + d p_0 - \delta(1 - p_0). \tag{S37}$$

Matching early-time rates,

$$\therefore r_{\mathrm{agg}}(0) = r_{\mathrm{bi}}(0) \quad \Rightarrow \quad f p_0^2 V_0 + d p_0 - \delta(1 - p_0) = p k_1 + (1 - p) k_2. \tag{S38}$$

With $\delta = k_2$ this becomes

$$\boxed{f p_0^2 V_0 + d p_0 - \delta(1 - p_0) = p k_1 + (1 - p) k_2.} \tag{S39}$$

Therefore for the PFU model, we have an exact analytical mapping for $\delta$ and have to numerically fit $d$, $f$, and $p_0$ as they are indeterminate from one other in the early-time regime (Equation S39).

##### Mathematical derivations of virus-host model steady states

We fix the total viral abundance from Equation 4,

$$V_1^* + V_2^* = V_{\text{target}}, \tag{S40}$$

where $V_1^*$ denotes free infectious virions, $V_2^*$ denotes non-infectious aggregates, and $V_{\text{target}}$ defines a reasonable standing abundance of total viruses, which we set to $5 \times 10^8$ virions$/L$. We also fix $n$, $d_{\text{A}}$, $\lambda$, $\omega$, and a reference infected fraction $q_0$ when there is no aggregation ($f = 0$), which we set to 0.01. Note that we set $g = 1$ because studies of viruses of microbes often use spatially resolved methods, including polony, VLP, and PFU assays, which count an aggregate as a single viral unit. We parameterize the disaggregation rate as $\delta = d_{\text{A}}/n$, where $n$ is a persistence factor that scales disaggregation relative to the single virion decay rate. This is consistent with the biphasic late time limit, for which $\delta = k_2$, where $k_2$ is the slower decay rate that governs the long tail of the biphasic decay curve (Equation S34). At high viral densities, aggregation can also contribute to the initial decay rate through the $fV_0$ term in Equation S39, yielding the corrected parameterization $\delta = (d_{\text{A}} + fV_0)/n$. However, incorporating this correction does not qualitatively alter the results in Figure 5. We therefore implement the simpler $\delta = d_{\text{A}}/n$ parameterization in Equation 4 to obtain the derivations below.

At steady state,

$$\dot{V}_2 = 0 = f(V_1^*)^2 - \frac{d_{\text{A}}}{n} V_2^*. \tag{S41}$$

$$V_2^* = \frac{nf}{d_{\text{A}}}(V_1^*)^2. \tag{S42}$$

Substituting this relationship into Equation S40 gives

$$V_{\text{target}} = V_1^* + \frac{nf}{d_{\text{A}}}(V_1^*)^2. \tag{S43}$$

$$\frac{nf}{d_{\text{A}}}(V_1^*)^2 + V_1^* - V_{\text{target}} = 0. \tag{S44}$$

$$V_1^* = \frac{-1 + \sqrt{1 + \frac{4nfV_{\text{target}}}{d_{\text{A}}}}}{2nf/d_{\text{A}}}. \tag{S45}$$

$$V_1^* = \frac{2V_{\text{target}}}{1 + \sqrt{1 + \frac{4nfV_{\text{target}}}{d_{\text{A}}}}}. \tag{S46}$$

We therefore define the free infectious virus fraction as

$$\boxed{z(f) = \frac{V_1^*}{V_{\text{target}}} = \frac{2}{1 + \sqrt{1 + \frac{4nfV_{\text{target}}}{d_{\text{A}}}}}.} \tag{S47}$$

and the non-infectious aggregate fraction as

$$\boxed{y(f) = 1 - z(f),} \tag{S48}$$

both of which we use to get the graphs seen in Figure 5a. Next, the infected host equation in Equation 4 is

$$\dot{I} = \phi_{\mathrm{A}} S V_1 - (\lambda + \omega) I. \tag{S49}$$

At steady state,

$$\phi_{\mathrm{A}} S^* V_1^* = (\lambda + \omega) I^*. \tag{S50}$$

$$\frac{I^*}{S^*} = \frac{\phi_{\mathrm{A}} V_1^*}{\lambda + \omega}. \tag{S51}$$

We define the steady state infected fraction as

$$q = \frac{I^*}{S^* + I^*}. \tag{S52}$$

$$q(S^* + I^*) = I^*, \tag{S53}$$

$$qS^* = I^*(1 - q). \tag{S54}$$

$$\frac{I^*}{S^*} = \frac{q}{1 - q}. \tag{S55}$$

Substituting into Equation S51 gets:

$$\frac{q}{1 - q} = \frac{\phi_{\mathrm{A}} V_1^*}{\lambda + \omega}. \tag{S56}$$

We define $q_0$ as the infected fraction in the reference case where aggregation is absent and the adsorption rate equals the no-aggregation reference rate:

$$f = 0, \qquad \phi_{\mathrm{A}} = \phi_{\mathrm{NA}}. \tag{S57}$$

When $f = 0$, all virions are free, so

$$V_1^* = V_{\mathrm{target}}. \tag{S58}$$

Equation S56 therefore becomes

$$\frac{q_0}{1-q_0} = \frac{\phi_{\mathrm{NA}} V_{\mathrm{target}}}{\lambda + \omega}. \tag{S59}$$

We define

$$x = \frac{\phi_{\mathrm{A}}}{\phi_{\mathrm{NA}}}, \tag{S60}$$

$$\therefore \phi_{\mathrm{A}} = x\phi_{\mathrm{NA}}. \tag{S61}$$

Using Equation S47 and Equation S61, Equation S56 becomes

$$\frac{q}{1-q} = \frac{x\phi_{\mathrm{NA}} z(f) V_{\mathrm{target}}}{\lambda + \omega}. \tag{S62}$$

Substituting into Equation S59,

$$\frac{q}{1-q} = xz(f)\frac{q_0}{1-q_0}. \tag{S63}$$

$$q(1-q_0) = xz(f)q_0(1-q). \tag{S64}$$

$$q(1-q_0) = xz(f)q_0 - xz(f)q_0 q. \tag{S65}$$

$$q(1-q_0) + xz(f)q_0 q = xz(f)q_0. \tag{S66}$$

$$q\left[1 - q_0 + xq_0 z(f)\right] = xq_0 z(f). \tag{S67}$$

$$q(f, x) = \frac{xq_0 z(f)}{1 - q_0 + xq_0 z(f)}. \tag{S68}$$

Substituting Equation S47 for $z(f)$ and expanding $x$ gives:

$$\boxed{q\left(f, \frac{\phi_{\mathrm{A}}}{\phi_{\mathrm{NA}}}\right) = \frac{2\left(\frac{\phi_{\mathrm{A}}}{\phi_{\mathrm{NA}}}\right) q_0}{(1-q_0)\left[1 + \sqrt{1 + \frac{4nfV_{\mathrm{target}}}{d_{\mathrm{A}}}}\right] + 2\left(\frac{\phi_{\mathrm{A}}}{\phi_{\mathrm{NA}}}\right) q_0}}. \tag{S69}$$

The percentage of infected hosts shown in Figure 5b is therefore

$$\boxed{100 \times q\left(f, \frac{\phi_{\mathrm{A}}}{\phi_{\mathrm{NA}}}\right)}. \tag{S70}$$

*Non-identifiability of adsorption and aggregation at fixed virus and infected prevelances*

We next ask whether different combinations of adsorption and aggregation can produce the same observable steady states. We therefore fix both the total viral abundance and steady state infected fraction,

$$V_1^* + V_2^* = V_{\text{target}}, \qquad \frac{I^*}{S^* + I^*} = q. \tag{S71}$$

We first determine the parameters of the no-aggregation model (Equation 3) that produce this prescribed equilibrium. As $\dot{I}^* = 0$:

$$\phi_{\text{NA}} S_{\text{NA}}^* V_{\text{target}} = (\lambda + \omega) I_{\text{NA}}^*. \tag{S72}$$

From the fixed infected fraction in Equation S71,

$$\frac{I_{\text{NA}}^*}{S_{\text{NA}}^*} = \frac{q}{1-q}. \tag{S73}$$

$$\therefore \phi_{\text{NA}} V_{\text{target}} = (\lambda + \omega)\frac{q}{1-q}, \tag{S74}$$

$$\phi_{\text{NA}} = \frac{(\lambda + \omega) q}{(1-q) V_{\text{target}}}. \tag{S75}$$

Now as $\dot{S}^* = 0$:

$$0 = r S_{\text{NA}}^* \left(1 - \frac{S_{\text{NA}}^* + I_{\text{NA}}^*}{K}\right) - \phi_{\text{NA}} S_{\text{NA}}^* V_{\text{target}} - \omega S_{\text{NA}}^*. \tag{S76}$$

Substituting Equation S74 gives

$$r\left(1 - \frac{S_{\text{NA}}^* + I_{\text{NA}}^*}{K}\right) = \omega + (\lambda + \omega)\frac{q}{1-q}. \tag{S77}$$

$$S_{\text{NA}}^* + I_{\text{NA}}^* = \frac{K}{r}\left[r - \omega - \frac{(\lambda+\omega)q}{1-q}\right]. \tag{S78}$$

Substituting Equation S73 gives

$$I_{\text{NA}}^* = \frac{Kq}{r}\left[r - \omega - \frac{(\lambda+\omega)q}{1-q}\right]. \tag{S79}$$

Now as $\dot{V}^* = 0$,

$$0 = \beta\lambda I^*_{\mathrm{NA}} - \phi_{\mathrm{NA}} S^*_{\mathrm{NA}} V_{\mathrm{target}} - d_{\mathrm{NA}} V_{\mathrm{target}}. \tag{S80}$$

Substituting Equation S72 gives

$$d_{\mathrm{NA}} V_{\mathrm{target}} = (\beta\lambda - \lambda - \omega) I^*_{\mathrm{NA}}. \tag{S81}$$

Substituting Equation S79 into Equation S81 gives

$$d_{\mathrm{NA}} = \frac{K(\beta\lambda - \lambda - \omega)q}{r V_{\mathrm{target}}} \left[ r - \omega - \frac{(\lambda + \omega)q}{1 - q} \right]. \tag{S82}$$

We now consider the aggregation model under the same constraints in Equation S71. $\dot{I}^* = 0$ gives

$$\phi_{\mathrm{A}} S^*_{\mathrm{A}} V^*_1 = (\lambda + \omega) I^*_{\mathrm{A}}. \tag{S83}$$

$$\frac{I^*_{\mathrm{A}}}{S^*_{\mathrm{A}}} = \frac{q}{1 - q}. \tag{S84}$$

$$\phi_{\mathrm{A}} V^*_1 = (\lambda + \omega) \frac{q}{1 - q}. \tag{S85}$$

Which equating to Equation S74 gives

$$\phi_{\mathrm{A}} V^*_1 = \phi_{\mathrm{NA}} V_{\mathrm{target}}. \tag{S86}$$

$$x = \frac{\phi_{\mathrm{A}}}{\phi_{\mathrm{NA}}}. \tag{S87}$$

$$V^*_1 = \frac{V_{\mathrm{target}}}{x}. \tag{S88}$$

$$V^*_2 = V_{\mathrm{target}} - V^*_1 = V_{\mathrm{target}} \left( 1 - \frac{1}{x} \right). \tag{S89}$$

We next determine the decay rate $d_{\mathrm{A}}$ required to maintain the same infected fraction in the aggregation model. As $\dot{S}^* = 0$,

$$0 = r S^*_{\mathrm{A}} \left( 1 - \frac{S^*_{\mathrm{A}} + I^*_{\mathrm{A}}}{K} \right) - \phi_{\mathrm{A}} S^*_{\mathrm{A}} V^*_1 - \omega S^*_{\mathrm{A}}. \tag{S90}$$

Substituting Equation S85 gives

$$S_A^* + I_A^* = \frac{K}{r}\left[r - \omega - \frac{(\lambda+\omega)q}{1-q}\right]. \tag{S91}$$

$$\therefore I_A^* = \frac{Kq}{r}\left[r - \omega - \frac{(\lambda+\omega)q}{1-q}\right]. \tag{S92}$$

Thus, comparison of Equation S92 and Equation S79 shows that fixing $q$ fixes the same steady state infected host abundance in both models,

$$I_A^* = I_{NA}^*. \tag{S93}$$

Setting $\dot{V_1}^* = \dot{V_2}^* = 0$ in the aggregation model (Equation 4) gives

$$0 = \beta\lambda I_A^* - 2f(V_1^*)^2 + 2\delta V_2^* - \phi_A S_A^* V_1^* - d_A V_1^*. \tag{S94}$$

$$f(V_1^*)^2 = \delta V_2^*. \tag{S95}$$

Substituting Equation S95 into Equation S94 gives

$$0 = \beta\lambda I_A^* - \phi_A S_A^* V_1^* - d_A V_1^*. \tag{S96}$$

Substituting Equation S83 into Equation S96 gives

$$d_A V_1^* = (\beta\lambda - \lambda - \omega) I_A^*. \tag{S97}$$

Equating to Equation S81 and using Equation S93 gives

$$d_A V_1^* = d_{NA} V_{\text{target}}. \tag{S98}$$

Substituting Equation S88 into Equation S98 gives

$$d_A \frac{V_{\text{target}}}{x} = d_{NA} V_{\text{target}}, \tag{S99}$$

$$\therefore d_A = x d_{NA}, \tag{S100}$$

Substituting $\delta = d_A/n$ into Equation S95 gives

$$f(V_1^*)^2 = \frac{d_A}{n} V_2^*, \tag{S101}$$

$$f = \frac{d_{\mathrm{A}} V_2^*}{n(V_1^*)^2}. \tag{S102}$$

Substituting Equation S100, Equation S88, and Equation S89 into Equation S102 gives

$$f = \frac{(x d_{\mathrm{NA}}) V_{\mathrm{target}} \left(1 - \frac{1}{x}\right)}{n \left(V_{\mathrm{target}}/x\right)^2}. \tag{S103}$$

$$f = \frac{d_{\mathrm{NA}}}{n V_{\mathrm{target}}} x^2 (x - 1). \tag{S104}$$

Finally, substituting $x = \phi_{\mathrm{A}}/\phi_{\mathrm{NA}}$ from Equation S87 gives

$$\boxed{f = \frac{d_{\mathrm{NA}}}{n V_{\mathrm{target}}} \left(\frac{\phi_{\mathrm{A}}}{\phi_{\mathrm{NA}}}\right)^2 \left(\frac{\phi_{\mathrm{A}}}{\phi_{\mathrm{NA}}} - 1\right).} \tag{S105}$$

Thus, Equation S105 defines combinations of adsorption and aggregation rates that preserve both the total viral abundance and infected host fraction, corresponding to Figure 5c.

##### Biphasic vs. exponential decay across datasets

The Boixereu et al (Guixa-Boixereu et al., 2002) dataset highlights the distinction between model preference and absolute fit quality, where although AICc did not strongly favor either model, the biphasic model fit the data far better than the exponential model by goodness of fit ($R^2_{bi} = 1.00$, $R^2_{exp} = 0.34$, Figure 1f). This likely reflects the small sample size ($n = 5$) and the corresponding AICc penalty, since $\mathrm{AICc} = \mathrm{AIC} + \frac{2k(k+1)}{n-k-1}$ strongly penalizes additional parameters when $n$ is small. For example, with $n = 5$, the AICc correction term $\frac{2k(k+1)}{n-k-1}$ is 24 for the three parameter biphasic model ($k = 3$), but only $\frac{4}{3}$ for the one parameter exponential model ($k = 1$), making the penalizing correction term for the biphasic model approximately 18 times larger. This highlights the need for more sampling in viral decay datasets to allow for resolution between decay models. In contrast, Tsai et al.(Tsai et al., 2022), Pleyer et al.(Pleyer et al., 2024), and Fischer et al. 2004(Fischer et al., 2004) were excluded from both the full 17-dataset comparison (Figure 1) and the well-fit model summary (Figure 2) as the AICc-winning model, or both models, failed to meet the $R^2 = 0.8$ fit-quality threshold (Tsai et al.(Tsai et al., 2022), Pleyer et al.(Pleyer et al., 2024), and Fischer et al. 2004(Fischer et al., 2004) fits are shown in Figure S1).
